# Chaperone Complexes From The Endoplasmic Reticulum (ER) And The Cytosol Inhibit wt-p53 By Activation The ER To Cytosol Signaling

**DOI:** 10.1101/2023.08.01.551134

**Authors:** Salam Dabsan, Gali Zur, Ayelet Gilad, Aeid Igbaria

## Abstract

The Endoplasmic Reticulum (ER) is an essential sensing organelle responsible for the folding and secretion of almost one-third of eukaryotic cells’ total proteins. The ER contains numerous enzymes and chaperones which assist in oxidative protein folding and other posttranslational modifications. However, environmental, chemical, and genetic insults often lead to protein misfolding in the ER, accumulating misfolded proteins, altering homeostasis, and causing ER stress. Recently, we reported a novel ER surveillance mechanism by which proteins from the secretory pathway are refluxed to the cytosol to relieve the ER of its content during stress. In cancer cells, the refluxed proteins gain new pro-survival functions, thereby increasing cancer cell fitness. We termed this phenomenon <u>ER</u> to <u>CY</u>tosol <u>S</u>ignaling (or “**ERCY”)**. In yeast, ERCYS is regulated by HLJ1 (an ER-resident tail-anchored HSP40 cochaperone). Here, we found that in mammalian cells, HLJ1 has five putative orthologs possessing J-domains facing the cytosol. Among those, DNAJB12 and DNAJB14 appear to be the most significant. Mechanistically, we found that DNAJB12 and DNAJB14 bind the cytosolic HSC70 and its cochaperone – SGTA - through their cytosolically localized J-domains to facilitate ER-protein reflux to the cytosol. Moreover, we found that DNAJB12 is necessary and sufficient to drive this phenomenon to increase AGR2 reflux and inhibit wt-p53 during ER stress. Thus, we concluded that targeting the DNAJB12/14-HSC70/SGTA axis is a promising strategy to inhibit ERCYS and impair cancer cell fitness.

## INTRODUCTION

The Endoplasmic Reticulum (ER) is an essential sensing organelle that provides multiple functions, including protein synthesis, folding, and secretion of almost one-third of all proteins in eukaryotic cells. In addition, the ER controls the cellular redox state, calcium storage, lipid metabolism, xenobiotics, and drug detoxification [1-4]. The ER also coordinates the stress pathways significantly involved in maintaining the crosstalk between the intra- and the extracellular environment [5, 6].

In addition, the ER is the entry point of proteins in the secretory pathway; these proteins are synthesized on the ER-associated ribosomes and destined to be secreted or targeted to the membrane[7-9]. In the ER, the nascent peptides gain their correctly folded structure and other posttranslational modifications through the activity of ER-resident chaperones. If correctly folded, those proteins exit the ER to their destination. Unfolded/misfolded proteins translocate to the cytosol, where they are ubiquitinylated and degraded by the proteasome in an ER-associated degradation (ERAD) mechanism[8, 10-13].

Accumulating misfolded proteins in the ER activates a signaling pathway called the Unfolded Protein Response (UPR) [14]. Initially, the UPR aims to regain homeostasis by increasing the ER folding capacity by making new ER chaperones. At the same time, the UPR decreases the number of substrates entering the ER through the activity of the inositol-requiring enzyme (IRE1) and protein kinase RNA-like ER kinase (PERK). Once activated, the IRE1 RNAse domain cleaves the mRNA of a transcriptional factor called Xbp1 to increase the chaperones in the ER[15-19]. Moreover, high IRE1 activity causes a massive degradation of mRNAs translated by ribosomes in proximity to the ER membrane. This regulated IRE1-dependent degradation (RIDD) mainly targets those mRNAs that primarily encode for proteins in the secretory pathway to decrease the protein load on the ER[20]. On the other hand, PERK activation results in eIF2α phosphorylation and inhibition of global protein translation[13, 21]. Thus, the ERAD and the UPR arms (PERK and IRE1) aim to decrease the number of client proteins in the ER, thereby reducing the ER protein load.

Along the same lines, we recently reported a novel ER surveillance mechanism by which proteins from the secretory pathway are refluxed from the ER to the cytosol to relieve the ER of its contents during stress. This conserved mechanism (from yeast to humans) targets a wide range of ER-resident and secretory proteins[22-24]. The reflux process is constitutively activated in cancer cells, causing many proteins to be enriched in the cytosolic fraction in cultured cancer cells, murine models of brain tumors, and human patients. In the cytosol, the refluxed proteins gain new pro-survival functions and thus increase cancer cell fitness. This was shown using the ER-resident PDI-like protein AGR2 that is refluxed from the ER of cancer cells to interact and inhibit the activity of pro-survival proteins in the cytosol, especially wt-p53. We named this phenomenon “ER to cytosol signaling” or (**ERCYS**)[23].

Our previous work in yeast showed that the reflux process depends on the activity of chaperones from both the ER and the cytosol and is mainly regulated by HLJ1 -an ER-resident tail-anchored HSP40 cochaperone[22, 24]. Upon ER stress, HLJ1 interacts with the cytosolic machinery to activate the removal of those proteins from the ER to the cytosol. In the cytosol, those refluxed proteins remain in protein complexes with cytosolic chaperones[22]. In this study, we found that HLJ1 is conserved through evolution and that mammalian cells have five putative functionality orthologs of the yeast HLJ1. Those five DNAJ-proteins (DNAJB12, DNAJB14, DNAJC14, DNAJC18, and DNAJC30) reside within the ER membrane with a J-domain facing the cytosol [25, 26]. Moreover, two putative orthologs, DNAJB12 and DNAJB14, are strongly related to the yeast HLJ1 and are sufficient and essential for determining cells’ fate during ER stress by regulating ERCYS. Their role in ERCYS and cells’ fate determination depends on their HPD motif in the J-domain. Downregulation of DNAJB12 and DNAJB14 increases cell toxicity and wt-p53 activity during etoposide treatment. Here, we propose a novel mechanism by which proteins are refluxed from the ER to the cytosol using chaperones and cochaperones from the ER and the cytosol to inhibit the activity of wt-p53 in cancer cells.

## RESULTS

### 1. Five Putative Orthologs of the Yeast HLJ1 in Mammalian Cells

Initially, we sought to identify factors that cause ER to cytosol reflux to affect wt-p53 activity in cancer cells. For this, we searched for orthologs of the yeast HLJ1 that we recently identified as an essential component in the reflux of ER proteins in S. cerevisiae[23]. Protein alignment of the yeast HLJ1p showed high amino acids similarity to the mammalian DNAJB12 and DNAJB14 with a very high score (Table S1 and Figure 1A). DNAJB12 and DNAJB14 are highly conserved in different species through evolution from yeast to humans (Figure 1A-1B and Table S1). Interestingly, other DNAJ proteins were also found in our list of hits. Among those, we could find DNAJC14 and DNAJC18, both recently identified as DNAJC-proteins that are expressed on the ER membrane with a similar topology as the yeast HLJ1, with a J-domain facing the cytosol (Figure 1A-C, Figure S1, and Table S1). An intensive analysis of the entire J-domain of DNAJB12, DNAJB14, DNAJC14, and DNAJC18 revealed highly conserved J-domains in humans and other species (Figure 1A and 1C). In addition, three amino acids, HPD, are also conserved in all the tested orthologs (Figure 1C-D and Figure S1). The HPD motif is conserved within the J-domain of those DNAJ proteins and is responsible for binding the cochaperones and the HSP70/HSC70 chaperones.

**Figure 1:**
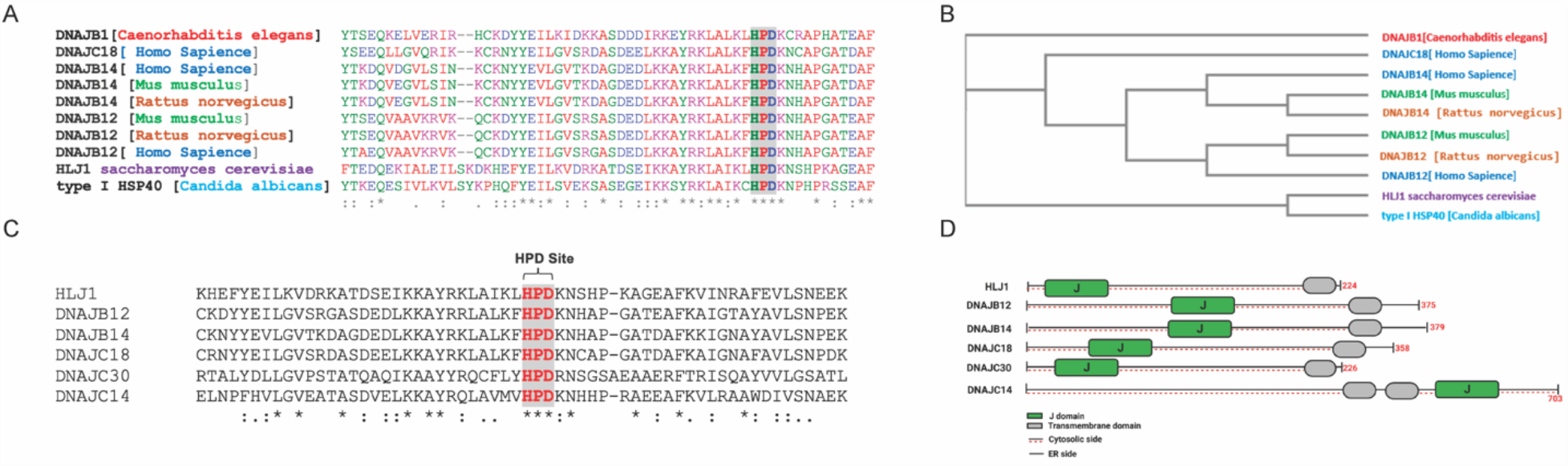
HLJ-1 is conserved from yeast to humans. **(A) alignment showing the** conservation of the yeast-HLJ1 and their HPD domain from yeast to human. (**B)** Phylogenic analysis showing the conservation of HLJ1 in different species. (**C)** the HPD motif within the J-domain is conserved in HLJ-1 and its putative human orthologs DNAJB12, DNAJB14, DNAJC14, DNAJC18, and DNAJC30. **(D)** Schematic showing the different domains of HLJ-1 and its orthologs, including a cytosolic J-domain, transmembrane domain, and Endoplasmic Reticulum (ER) domain.

Finally, it is important to emphasize that the four identified putative orthologs and another DNAJC protein - DNAJC30 - were shown to localize to the ER membrane with a J-domain facing the cytosol [25, 26]. DNAJC30 shares the same topology on the ER membrane, consisting of one transmembrane domain (or more for the DNAJC30) and polar residues (Figure 1D and Figure S1). These data suggest that mammalian cells may have developed several orthologs of the yeast HLJ1. The high similarity of DNAJB12 and DNAJB14 to the yeast HLJ-1 indicates that they (DNAJB12 and 14) may be the ones that carry most of the yeast HLJ1 functions. In addition, DNAJC14, DNAJC18, and DNAJC30 are putative orthologs to the yeast HLJ1, based on their topology and J-domain, but with lower similarity scores.

### 2. DNAJB12 AND DNAJB14 Regulate AGR2 Reflux from the ER to the Cytosol

Previously we showed that, during ER stress, the ER resident anterior gradient 2 (AGR2), a protein disulfide isomerase family member, and other ER-resident proteins are refluxed from the ER to the cytosol to interact and inhibit wt-p53 in different cancer cell lines [23]. Here, we wanted to test the role of our two strongest hits, DNAJB12 and DNAJB14, in the reflux of ER-resident proteins and wt-p53 activity. For this, we transfected A549 cells with shRNA against DNAJB12, DNAJB14, or both at different time points (Fig S2A-B). We treated the J-protein-silenced A549 cells with two different ER stress inducers: thapsigargin (Tg) -a sarco-ER Ca2+ ATPase, and tunicamycin (Tm) -N-linked-glycosylation inhibitor. In the WT and J-protein-silenced A549 cells, there were no differences in the cytosolic enrichment of the three ER resident proteins AGR2, DNAJB11, and HYOU1 in normal and unstressed conditions(Figure 2A-C and Figure S2C). Tm and Tg treatment increased the cytosolic enrichment of AGR2, DNAJB11, and HYOU1, mainly in the WT cells. DNAJB12-silenced cells are slightly affected in AGR2 and DNAJB11 cytosolic accumulation but not HYOU1. On the other hand, DNAJB14-silenced cells were only affected in AGR2 but not in DNAJB11 or HYOU1 cytosolic accumulation. Despite the slight decrease in the cytosolic accumulation of the selected proteins in the single knockouts, some of those changes were statistically significant (Figure 2A-C and Figure S2C).

**Figure 2:**
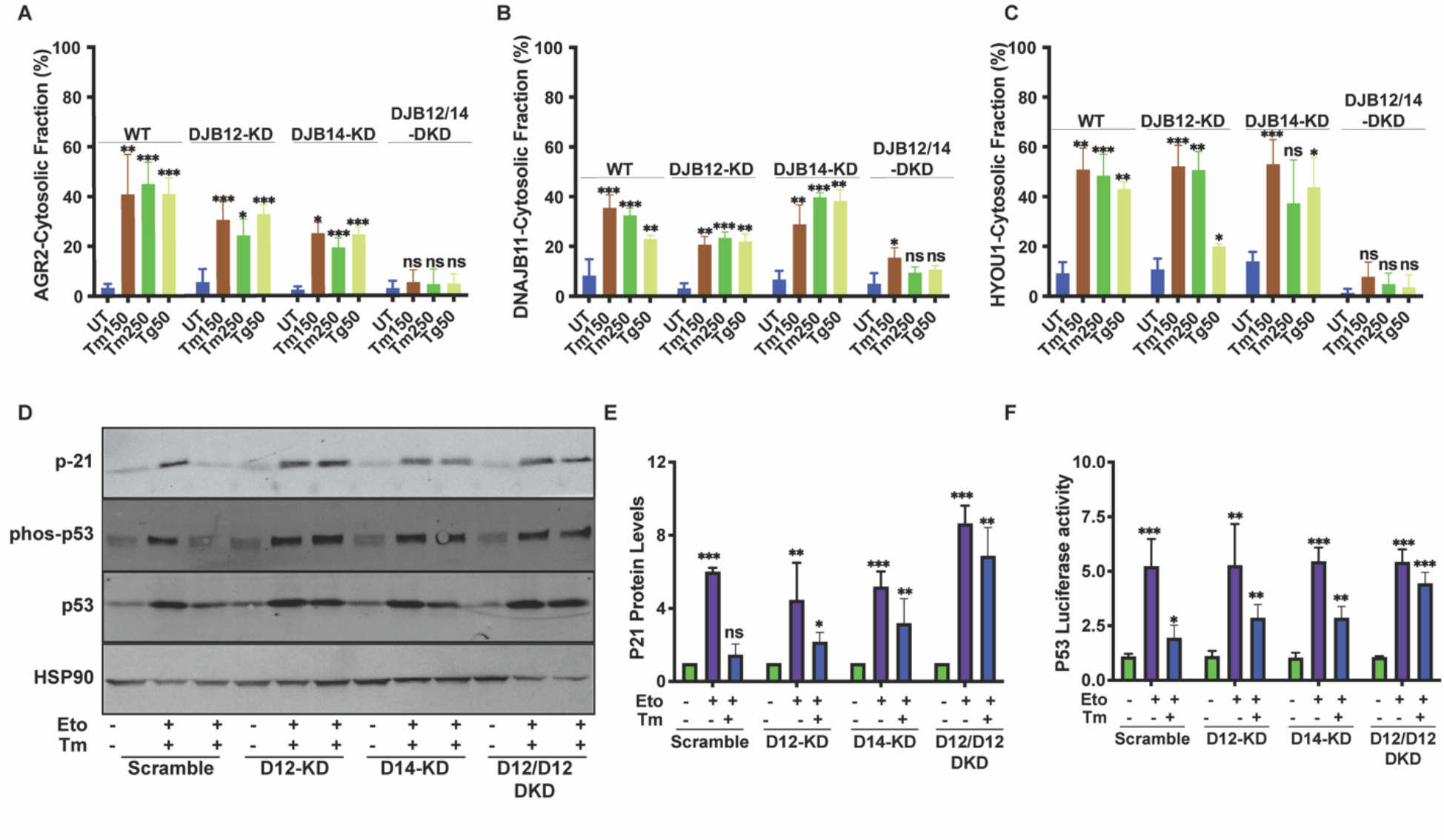
DNAJB12 and DNAJB14 are necessary for AGR2 reflux and wt-p53 inhibition in cancer cells. **(A-C)** Quantification of the subcellular protein fractionation (Digitonin fraction) of AGR2, DNAJB11, and HYOU1 in A549 cells as shown in (Figure S2C). **(D)** Representative immunoblot of P-21, phospho-p53, total-p53 (DO-1), and HSP90 in control cells and cells lacking DNAJB12, DNAJB14, or both. **(E)** Quantification of P-21 levels as shown in D. **(F)** A549 were transfected with scramble siRNA (scramble), DNAJB12-targeted siRNA (D12-KD), DNAJB14-targeted siRNA (D14-KD) or both. After 24 hours, cells were transfected with p53-luciferase construct. Cells were treated with etoposide for 2 hours to induce wt-p53, followed by tunicamycin treatment for 16 hours, and luciferase experiments were performed. Biological triplicates, mean ± SD calculated using Prism 9 (GraphPad). (***p<0,001, **p<0,01, *<0.05).

Because DNAJB12 and DNAJB14 are highly homologous and probably have overlapping functions, we tested the ER protein reflux in the double knockdown of DNAJB12 and DNAJB14 under the same conditions of ER stress. During stress, DNAJB12/DNAJB14 double knockdown was highly affected in the cytosolic accumulation of all three tested proteins (Figure 2A-C and Figure S2C). Those data clearly show that the reflux of AGR2 and other ER-resident proteins depends on the presence of both DNAJB12 and DNAJB14. These data also further confirm our hypothesis that the DNAJB12 and DNAJB14 are strong hits of the yeast HLJ1 and are necessary to drive ER-cytosol reflux.

We then wanted to examine whether the gain of function of AGR2 and the inhibition of wt-p53 depends on the activity of DNAJB12 and DNJAB14. We assayed the phosphorylation state of wt-p53 and p21 protein expression levels (a downstream target of p53 signaling) during etoposide treatment. Tm addition to cells treated with etoposide resulted in a reduction in wt-p53 phosphorylation, and as a consequence, the p21 protein levels were also decreased (Figure 2D-E). Cells lacking DNAJB12 and DNAJB14 have partial protection in wt-p53 phosphorylation and p21 protein levels. Silencing both proteins in A549 cells increased wt-p53 phosphorylation and p21 levels (Figure 2D-E). Moreover, similar results were obtained when we assayed the transcriptional activity of wt-p53 in cells transfected with a luciferase reporter under the p53-DNA binding site (Figure 2F). These data confirm that DNAJB12 and DNAJB14 are involved in ER protein reflux and the inhibition of wt-p53 activity during ER stress.

Interestingly, when we overexpressed WT-DNAJB12 in A549 cells, it was sufficient to decrease wt-p53 luciferase activity in cells treated with etoposide compared to the control cells (Figure 3A). To test whether this decrease in wt-p53 activity requires a functional DNAJ protein, we generated mutations in the HPD motif in the J-domain -needed for its cochaperone activity and the recruitment of HSC70-by substituting the HPD to QPD at position 139 or 136 for DNAJB12 and DNAJB14, respectively. Although the WT and the QPD mutants were expressed at the same levels (Figure S3A-B), overexpression of the DNAJB12-QPD mutant was incapable of decreasing the p53-luciferase activity under the same conditions (Figure 3A). On the other hand, overexpression of DNAJB14 alone or the DNAJB14-QPD mutant could not inhibit the luciferase activity of wt-p53 (Figure 3B). Those data indicate that DNAJB12 is sufficient to inhibit p53 activity in A549 cells treated with etoposide in a mechanism that requires its HPD motif and a functional J-domain.

**Figure 3:**
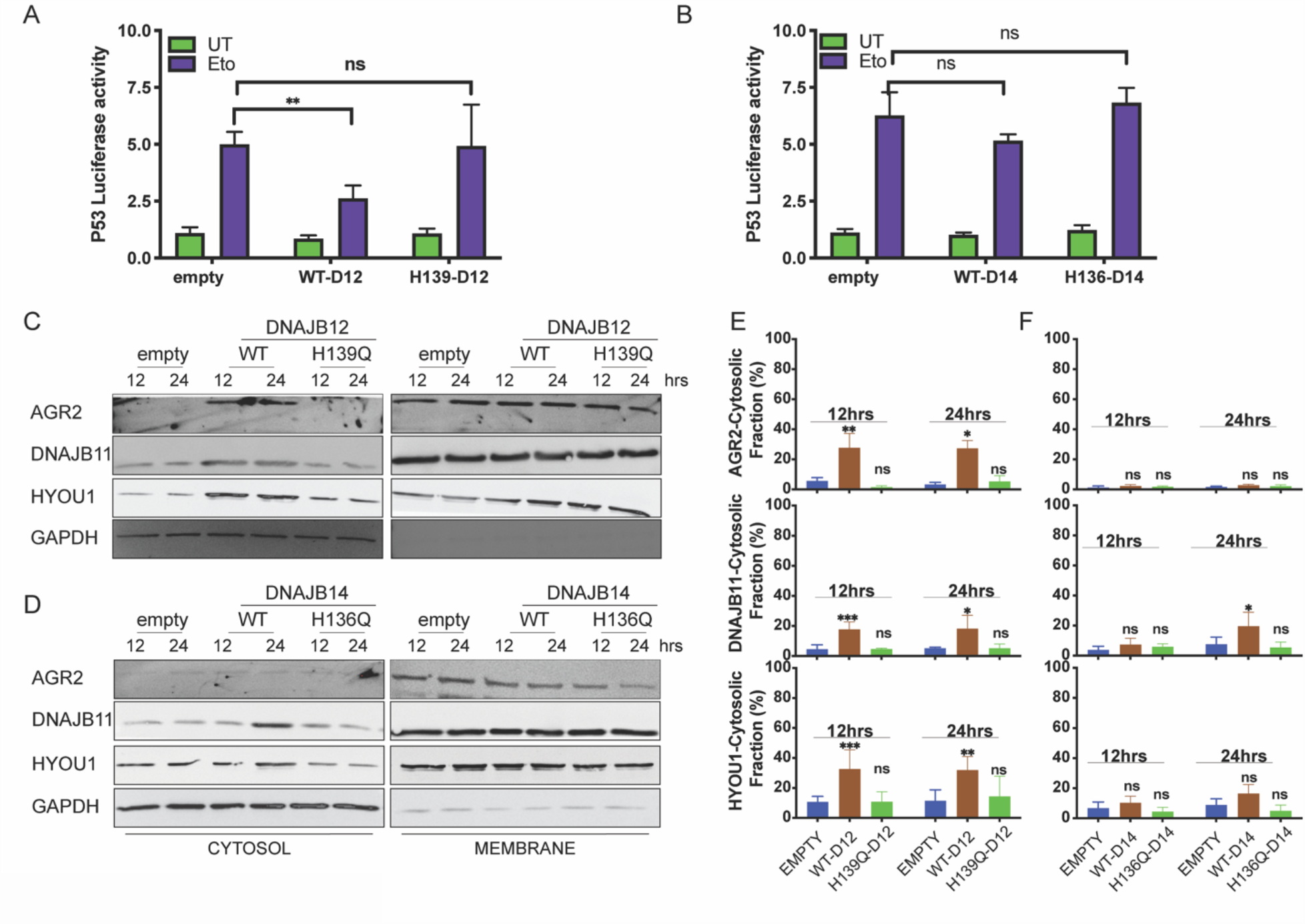
DNAJB12 but not DNAJB14 is sufficient to expel AGR2 and other ER resident proteins to inhibit wt-p53 activity. **(A-B) Cells were cotransfected with pCDNA3 plasmid expressing DNAJB12, DNAJB14, or the empty plasmid and** p53-luciferase construct, after 24-hour cells were treated with etoposide for 4 hours to induce wt-p53 and luciferase experiments were performed. Biological triplicates, mean ± SD calculated using Prism 9 (GraphPad). (***p<0,001, **p<0,01, *<0.05). **(C-D)** Subcellular protein fractionation (Digitonin fraction) of AGR2, DNAJB11, and HYOU1in cells overexpressing WT and QPD mutant of DNAJB12 and DNAJB14, respectively, at different time points. **(E-F)** Quantification of the subcellular protein fractionation of AGR2, DNAJB11, and HYOU1 in A549 cells as shown in (C-D).

We previously found that the yeast HLJ1 is sufficient to induce the reflux of ER proteins from the ER to the cytosol [22]. Thus, we hypothesized that overexpression of DNAJB12 may induce ERCYS and result in wt-p53 inhibition (figure 3A-B). To test this hypothesis, we assayed the reflux of AGR2 during ERCYS. First, we tested whether DNAJB12 and DNAJB14 are sufficient to promote ERCYS in cancer cells without ER stress and just by mass action. We performed subcellular protein fractionation in cells overexpressing DNAJB12 or DNAJB14 and their QPD mutants. Overexpressing DNAJB12 in A549 cells was sufficient to cause reflux of AGR2, DNAJB11, and HYOU1 at 12 and 24 hours post-induction (Figure 3C and 3E). On the other hand, overexpressing DNAJB14 alone was insufficient to increase the cytosolic accumulation of the tested ER-resident proteins, compared to DNAJB12 overexpression (Figure 3D and 3F). Those data confirm that DNAJB12 is sufficient to induce ERCYS and the reflux of AGR2 to the cytosol to inhibit wt-p53 activity. This reflux of ER proteins depends on the activity of the J-domain of DNAJB12 and is independent of the UPR activation in cells overexpressing DNAJB12 (Figure S3D).

It is important to note that overexpression of DNAJB12 for more than 24 hours was toxic compared to cells overexpressing the DNAJB12-QPD mutant or the empty vector in A549 and T-REx-293 cells (Figure S3C). These data further indicate that DNAJB12 and DNAJB14 are true orthologs of the yeast HLJ1 (**H**igh-copy **L**ethal **J**-protein or HLJ1), which was also found to be toxic when overexpressed.

Overexpressing the DNAJB12-WT in A549 cells treated with tunicamycin maintained the reflux of AGR2, DNAJB11, and HYOU1. As expected, overexpressing DNAJB12-HPQ has a dominant negative effect on the ER-protein reflux during ER stress (Figure 4A-E). Interestingly, overexpressing DNAJB14-wt and DNAJB14-HPQ inhibited ER protein reflux during ER stress (Figure 4F-J). This latter result further strengthens the idea that a functional HPD motif of DNAJB12 is necessary for this phenomenon and that a proper ratio between DNAJB12 and DNAJB14 is needed. This suggests that the two proteins may have different functions when overexpressed, despite their overlapping and redundant functions.

**Figure 4:**
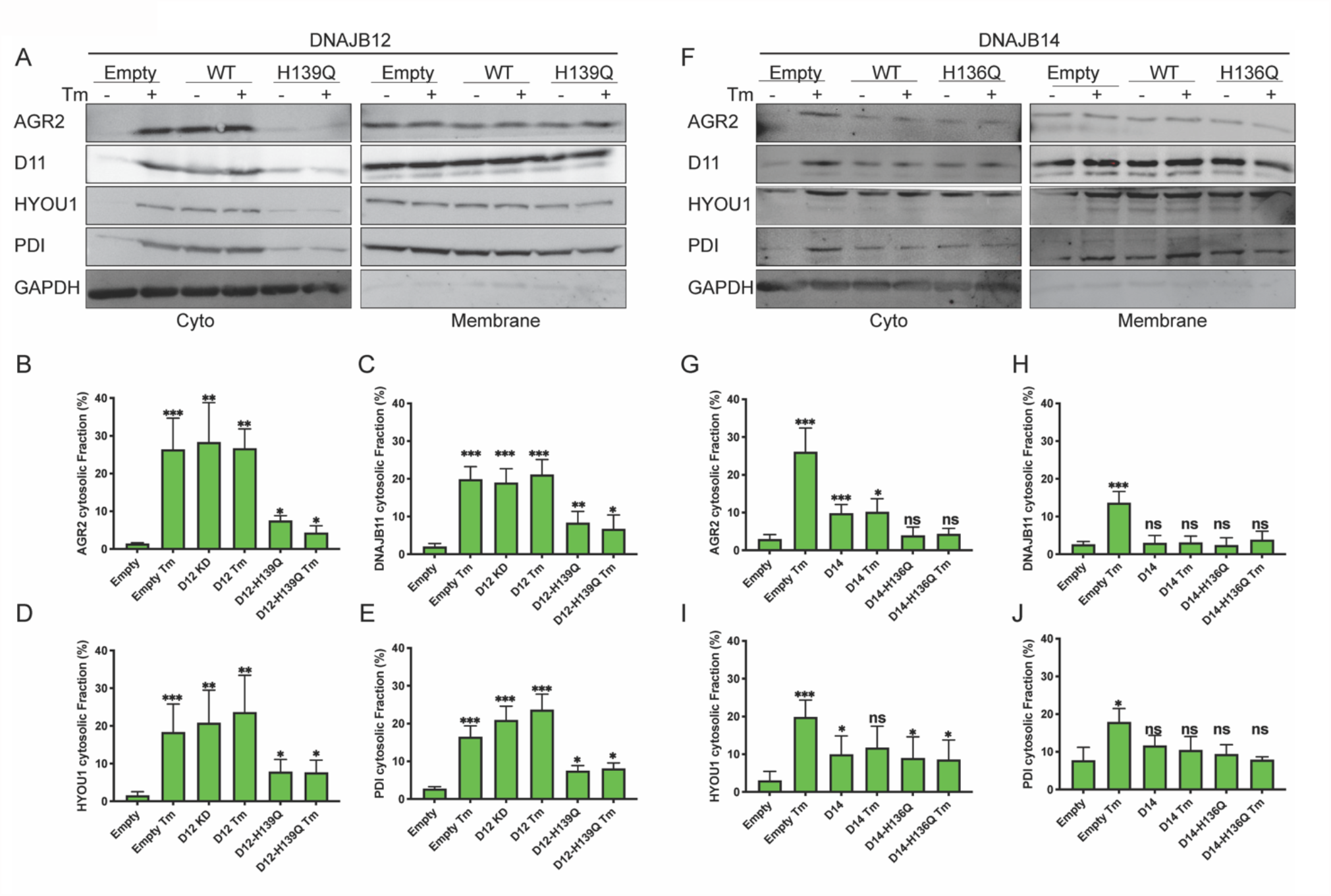
Functional DNAJB12 and 14 are necessary for ER protein reflux during ER stress. **(A)** Subcellular protein fractionation (Digitonin fraction) of AGR2, DNAJB11, and HYOU1in cells overexpressing either WT and QPD mutant of DNAJB12 during ER stress with tunicamycin (Tm). **(B-E)** Quantification of the subcellular protein fractionation of AGR2, DNAJB11, and HYOU1 in A549 cells as shown in A. **(F)** Subcellular protein fractionation (Digitonin fraction) of AGR2, DNAJB11, and HYOU1 in cells overexpressing WT and QPD mutant of DNAJB14 during ER stress with tunicamycin (Tm). **(G-J)** Quantification of the subcellular protein fractionation of AGR2, DNAJB11 and HYOU1 in A549 cells as shown in F. (***p<0,001, **p<0,01, *<0.05).

### 3. HSC70-SGTA Axis Regulates ER-Protein Reflux

Because the reflux process requires a functional HPD motif in the J-domain, we hypothesize that HSC70 and other cytosolic partners of ER-localized J-proteins may be necessary. To test this hypothesis, we performed coimmunoprecipitation experiments to assay the binding of DNAJB12/14, HSC70/SGTA (small glutamine-rich tetratricopeptide repeat, containing protein-α and cochaperone of HSC70), and the ER-localized proteins that are refluxed during stress. To assay the binding of DNAJB12 and DNAJB14 to the cytosolic partners, we used the FLAG-tagged DNAJB12 and the HA-tagged DNAJB14 and made stable lines using the Flp-In T-REx-293 cell lines (Thermofisher) for doxycycline-inducible gene expression. Overexpression of DNAJB12 in T-REx-293 cells shows an interaction between the HSC70 cochaperone (SGTA) and DNAJB12 in cells treated with doxycycline or doxycycline and tunicamycin (Figure 5A).

**Figure 5:**
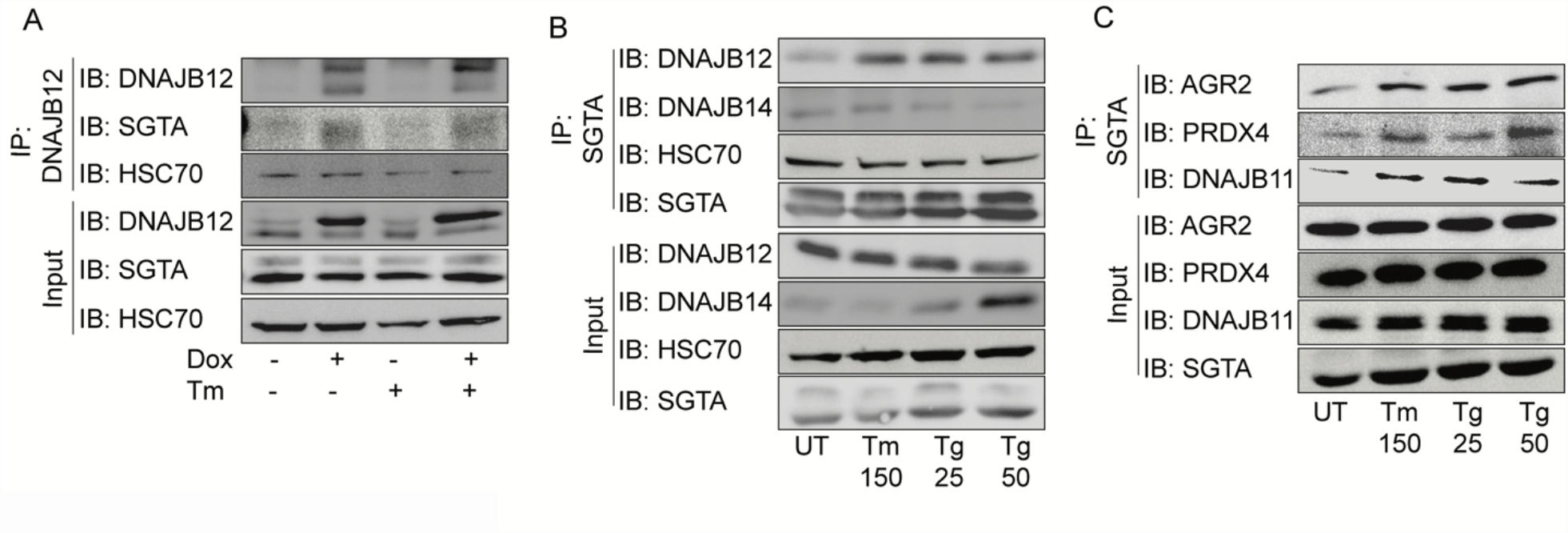
The cytosolic cochaperone SGTA is recruited by DAJB12 and interacts with the refluxed ER protein in the cytosol. **(A)** T-Rex cell lines overexpressing DNAJB12 using the Tet-On system for Doxycycline-inducible gene expression **were treated with doxycycline to induce DNAJB12 expression**. DNAJB12 interaction with SGAT was analyzed by COimmunoprecipitation assay. (B) Representative immunoblot showing the interaction between SGTA in one hand and DNAJB12, DNAJB14, and HSC70 in A549 cells treated with tunicamycin and thapsigargin as indicated. (C) Representative immunoblot showing the interaction between SGTA in one hand and the ER-resident proteins AGR2, PRDX4, and DNAJB11 in A549 cells treated with tunicamycin and thapsigargin as indicated.

We then tested whether the SGTA binds endogenous DNAJB12 and DNAJB14 in A549 cells treated with ER stress inducers. For this, we Immunoprecipitated SGTA and found that it binds DNAJB14 under normal and ER stress conditions. SGTA-DNAJB12 binding was enriched under ER stress compared to the control (Figure 5B). Finally and most importantly, SGTA immunoprecipitation showed an increased interaction between the cytosolic SGTA and the ER-resident proteins AGR2, PRDX4, and DNAJB11 (Figure 5C). These data are important because SGTA and the ER-resident proteins (PRDX4, AGR2, and DNAJB11) are known to be expressed in different compartments, and the interaction occurs only when those ER-resident proteins localize to the cytosol. Overall, those data show that the mechanism of action in mammalian cells is not different from that of yeast, where HLJ1 recruits cytosolic chaperones and cofactors to cause reflux. In mammalian cells, the reflux process depends on cochaperones from the ER membrane DNAJB12 and DNAJB14 and their J-domain facing the cytosol. During ERCYS, HSC70 and SGTA are recruited by DNAJB12 to facilitate the reflux of proteins to the cytosol.

### 4. The J Domains of DNAJB12 And DNAJB14 Regulate Cell Fate During ER Stress

Finally, we performed an XTT assay to measure cellular metabolic activity and to test cell viability and proliferation to show the physiological relevance of DNAJB12 and DNAJB14 inhibition on the proliferation of cancer cells. In control cells, etoposide decreased the proliferation index of A549 cells by almost two folds. Treating the cells with sub-toxic concentrations of tunicamycin decreased the toxicity caused by etoposide (Figure 6A). Silencing DNAJB12/DNAJB14 after etoposide treatment abolished the effect of Tm and resulted in no rescue of the proliferation of the double-knockout cells (Figure 6A). We then tested the role of SGTA on cell survival in cells treated with etoposide during ER stress. For this, we downregulated SGTA using the CRISPRi method reported earlier in [27] and tested the XTT assay in SGTA-silenced A549 cells. Etoposide treatment decreased the proliferation of A549 cells; a subtoxic concentration of Tm rescued this toxicity and increased the proliferation index (Figure 6B). In SGTA-silenced cells, subtoxic concentrations of Tm could not recover proliferation in a similar way seen with DNAJB12/DNAJB14 double knockout (Figure 6B). These data indicate that DNAJB12 and DNAJB14 protect cells from toxicity by refluxing ER proteins to the cytosol, where they gain new functions to inhibit p53 activity. These findings are significant and propose DNAJB12 and DNAJB14 as promising targets in restoring p53 activity and cancer cell sensitization to anti-cancer therapies. Moreover, these data suggest that SGTA is not only needed for the reflux of the ER proteins but also affects the wt-p53 activity in cancer cells treated with ER stress and etoposide (Figure 6C-D).

**Figure 6:**
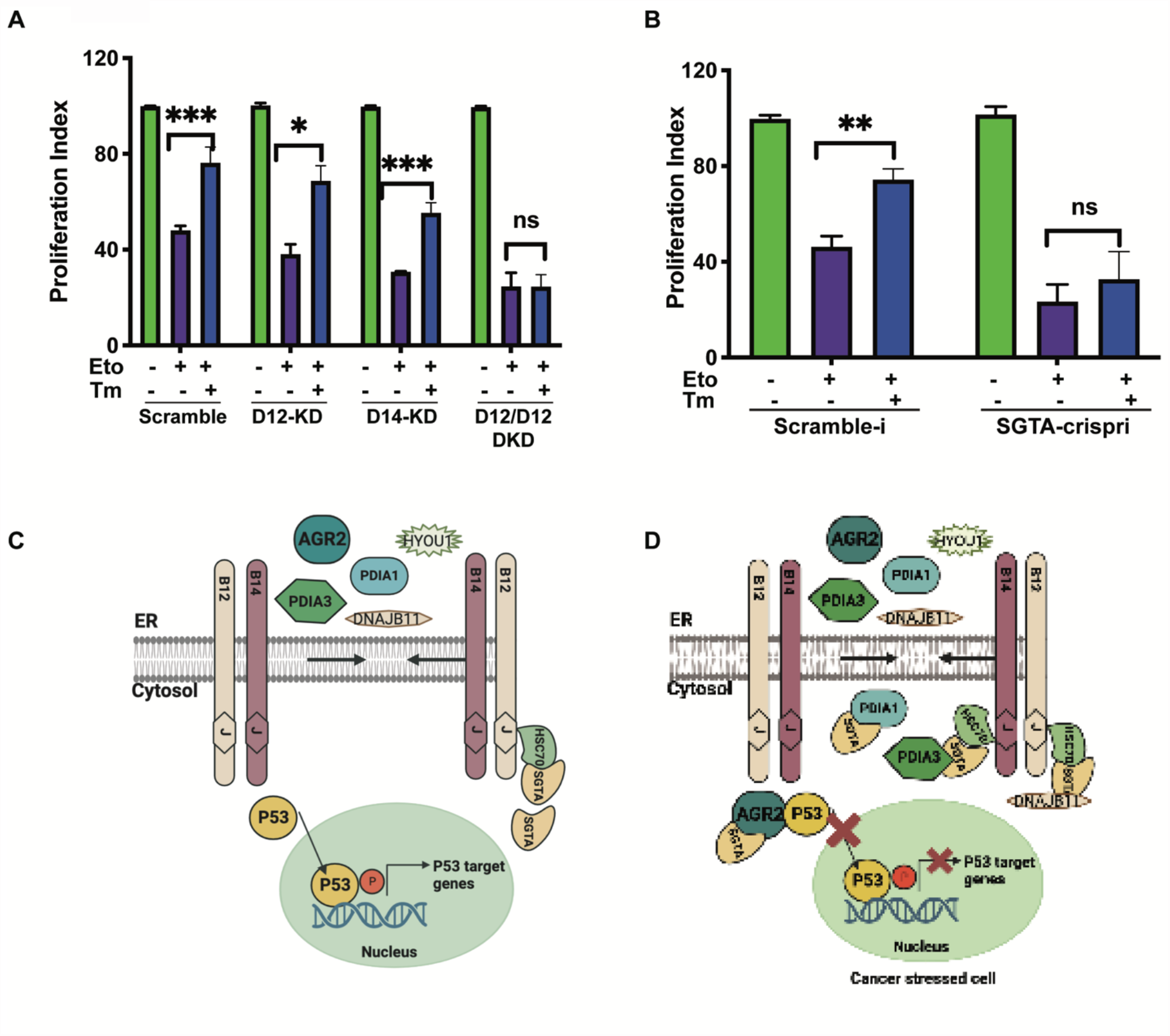
Silencing DNAJB12 and DNAJB14 or SGTA inhibits the rescue cell proliferation under subtoxic ER stress conditions. **(A)** XTT assay in A549 transfected with scramble siRNA (scramble), DNAJB12-targeted siRNA (D12-KD), DNAJB14-targeted siRNA (D14-KD), or both. After 24 hours, cells were treated with etoposide in the presence or absence of Tm for 48 hours. XTT assay in SGTA-silenced A549 cells using CRISPRi and treated with etoposide in the presence or absence of tunicamycin for 48 hours. Biological triplicates, mean ± SD calculated using Prism 9 (GraphPad). (***p<0,001, **p<0,01, *<0.05). **(C-D)** Cartoon showing our working model in cancer cells.in normal conditions, there is almost full compartmentation between the ER and the cytosol (C). Under ER stress conditions, DNAJB12 and DNAJB14 recruit cytosolic chaperones and cochaperones (SGTA) to expel AGR2 and other ER resident proteins to the cytosol and inhibit wt-p53 activity (D).

## DISCUSSION

ERCYS is a novel mechanism that acts as a pro-survival process to increase cancer cell fitness [23]. This is done by (1) decreasing ER-protein load and (2) refluxing of pro-survival proteins (such as AGR2) to gain new functions in the cytosol and inhibit proapoptotic protein signaling [23]. Despite that, the mechanism by which this process causes the reflux of those proteins to the cytosol is still unknown. Previously, we showed that the yeast cochaperone HLJ1 (ER-membrane localize Type-II HSP40 protein) is responsible for this phenomenon [22].

Our results show that ERCYS is regulated by the action of at least two ER-localized DNAJ proteins, namely DNAJB12 and DNAJB14 (Figure 1, Table S1, and Figure S1). Here, we bring evidences that DNAJB12 and DNAJB14 are the real orthologs of the yeast HLJ-1 by: (1) showing that DNAJB12 and DNAJB14 have a high similarity protein sequence with HLJ1, (2) both DNAJB12 and DNAJB14, on the one hand, and HLJ1, on the other hand, localize on the ER membrane with a J-domain facing the cytosol, (3) DNAJB12 and DNAJB14 were reported to facilitate the escape of nonenveloped viruses from the ER to the cytosol in a mechanism that involves chaperones from both the ER and the cytosol (in a similar manner to that of the role of HLJ1 in refluxing ER proteins to the cytosol), and (4) similarly to HLJ1, DNJAB12 is required for the degradation of a misfolded membranal protein - a role that was also reported for HLJ1. In addition, we propose that DNAJC14, DNAJC18, and DNAJC30 are homologs of DNAJB12 and DNAJB14 having an ER transmembrane domain and a J-domain facing the cytosol. Moreover, while in yeast, there is only one DNAJ protein on the ER membrane facing the cytosol, it is interesting to further study the role of the five orthologs in mammals and whether they have specificity towards different substrates and may reflux different sets of proteins. In addition, we found that DNAJB12 is toxic when overexpressed, similar to the yeast HLJ1. This toxicity could result from upregulated ERCYS activity that, at later time points, causes cell death by depleting the ER of its content.

We also showed that DNAJB12 and DNAJB14 are highly homologous and may have overlapping functions. Downregulation of either DNAJB proteins was partially necessary for ER protein reflux, but when both DNAJBs were downregulated, ERCYS was inhibited (Figure 2 and Figure S2). Moreover, DNAJB12 was sufficient to promote this phenomenon and cause ER protein reflux by mass action without causing ER stress (Figure 3, Figure 4, and Figure S3). Those data are very similar to the function of yeast HLJ1, being essential and sufficient for protein reflux. [22]. DNAJB12 overexpression was also sufficient to promote ERCYS by refluxing AGR2 and inhibit wt-p53 signaling in cells treated with etoposide (Figure 3). Moreover, the downregulation of DNAJB12 and DNAJB14 rescued the inhibition of wt-p53 during ER stress (Figure 2). These important observations indicate that wt-p53 inhibition is independent of ER stress and UPR activation but depends on the reflux of ER proteins to the cytosol.

Finally, ER stress is constitutively active in different cancer types, including those with wt-p53. AGR2 expression was also associated with the downregulation of wt-p53 activity. Moreover, high levels of wt-p53 activity were associated with higher AGR2 expression compared to tumors with mutant p53 [28, 29]. Here we show a new connection between those two signaling pathways. ERCYS is a novel mode of connection between ER stress and wt-p53 inhibition. In this connection, DNAJB12 and DNAJB14 are the link that regulates ER protein expulsion and wt-p53 activity. Our working model proposes that in non-cancer and resting cells, there is a high degree of segregation and separation between the content of the ER and the cytosol. In cancer cells, DNAJB12 and DNAJB14 oligomerize and recruit cytosolic chaperones and cochaperones (HSC70 and SGTA) to reflux AGR2 and other ER-resident proteins and to inhibit wt-p53 and probably different proapoptotic signaling pathways (Figure 5, and Figure 6C-6D). This mechanism is highly similar to the process suggested earlier regarding the escape of nonenveloped viruses[30]. This further strengthens our hypothesis that viruses have hijacked this evolutionarily conserved mechanism, leading to the escape of ER proteins to the cytosol.

## MATERIALS AND METHODS

### Cell culture and reagents

Human T-REx-293 and A549 cells were cultured in Dulbecco’ss modified eagle’s medium (DMEM) supplemented with 10% FBS and 1% antibiotics at 37°C in a 5% CO2 incubator. Tunicamycin (Tm) was purchased from Calbiochem. Thapsigargin (Tg), and Etoposide (Eto), were purchased from Sigma-Aldrich.

### Plasmids transfection

cDNA from A549 cells was used to PCR tag DNAJB12 and DNAJB14 with different tags. The PCR product was then digested and cloned into pCDNA5-FRT/To or pCDNA3(+) plasmids and transfected to A549 cells and T-REx-293 cells. The HPD to HPQ mutants were generated using Quickchange site-directed mutagenesis kit. FLAG-DNAJB12, HA-DNAJB14, and PG13-luc (Addgene plasmid # 16442) were transfected using lipofectamine 2000 (Thermo-Fisher Scientific) according to the manufacturer’s protocol.

### Western blots and antibodies used

Cells were washed with PBS (ice-cold), and whole cell lysates were collected using RIPA buffer (25 mM Tris/HCl pH 7.5, 150 mM NaCl, 1% sodium deoxycholate, 0.1% SDS, and 1% NP-40). Lysates were centrifuged at four °C for 10 minutes at 11000Xg, and proteins were quantified using BSA gold (Thermofisher) and loaded on SDS/PAGE. Primary antibodies (1:1000 diluted) were incubated overnight at 4°C. Fluorescent secondary antibodies were diluted 1:10000 and incubated for 1 hour before scanning using iBright Imaging Systems (Thermofisher). The list of antibodies is found in Table S3.

### Subcellular protein fractionation protocol

The fractionation protocol was used as described earlier [31]. In Brief, after a quick wash with ice-cold PBS, cells were trypsinized for 5 minutes and pelleted at 100RCF for 5 minutes at 4°C. The pellets were washed with ice-cold PBS, resuspended in buffer-1 (50 mM HEPES pH 7.4, 150 mM NaCl, 10 μg/ml digitonin (25ug/mL)), and incubated for 10 minutes at 4°C with rotation. Cells were pelleted at 2000RCF for 5 minutes at 4°C, and the cytosolic fraction (supernatant) was collected. The pellet was dissolved in Buffer-2 (50 mM HEPES pH 7.4, 150 mM NaCl, 1% NP40) and incubated for 30 min on ice. The cells were centrifuged at 7000RCF for 5 min at 4°C, and the supernatants were collected (membranal fraction).

### Coimmunoprecipitation protocol

Cells were grown as indicated for 24 hours. Tunicamycin and thapsigargin were added for 16 hours. Cells were then collected using Co-IP buffer (50 mM Tris/HCl pH 8, 150 mM NaCl, 0.5% TritonX100, and 1 mM EDTA) and left for 30 min on ice. The lysates were then centrifuged for 10 minutes at 11000Xg at 4°C. Proteins were quantified using BSA gold (Thermofisher), an equal amount of proteins were taken for each IP, and primary antibodies (1 μg Ab/1000 μg protein) were incubated overnight at 4°C. Washed Dynabeads protein A (Life Technologies) were mixed with the protein/antibodies mixture and incubated for 3 hours at 4°C with gentle rotation. After three washes with Co-IP buffer, the beads were transferred to a clean tube, eluted with 50 μl of Laemmli sample buffer, heated for 5 min at 100°C, and loaded to SDS/PAGE.

### RNA isolation and Real-Time PCR (qPCR)

RNA was isolated from T-REx-293 and A549 cells using the NucleoSpin RNA Mini kit for RNA purification (Macherey-Nagel). 500 ng of total RNA was used for cDNA synthesis using Maxima™ Reverse Transcriptase (ThermoFisher). Syber green (Thermofisher) was used for the qPCR reaction using (Quantstudio). Relative mRNA levels and gene expression levels were normalized to GAPDH or HPRT1. Primer sequences were used as described in [32].

### Cell Proliferation assay

The XTT cell proliferation kit (BioInd, SARTORIUS) was performed according to the manufactural protocol. 2500 cells were grown in 100μL in 96 wells and incubated for 24-96hours in 5% CO2 incubator at 37°C with different stressors. 50μL of XTT reagent solution/activation mix was added to each well and incubated for 4 hours at 37°C. Absorbance was then measured at 450 and 690 nanometers.

## CONFLICT OF INTEREST

The authors declare that they have no conflict of interest.

## AUTHOR CONTRIBUTIONS

SD, Gz, and AI designed the experiments. SD performed the experiments with GZ and AG. AI conceived the project, supervised the research, and wrote the manuscript with intellectual input and editing from all authors.

## Fundings

This work was supported by the Israel Science Foundation (ISF, grant No. 977/21)

## Notes

### Competing Interest Statement

The authors have declared no competing interest.

## REFERENCES

1. Reid, D.W. and C.V. Nicchitta, Diversity and selectivity in mRNA translation on the endoplasmic reticulum. Nat Rev Mol Cell Biol, 2015. 16(4): p. 221–31.

2. Rapoport, T.A., Protein translocation across the eukaryotic endoplasmic reticulum and bacterial plasma membranes. Nature, 2007. 450(7170): p. 663–9.

3. Jan, C.H., C.C. Williams, and J.S. Weissman, Principles of ER cotranslational translocation revealed by proximity-specific ribosome profiling. Science, 2014. 346(6210): p. 1257521.

4. Goyal, U. and C. Blackstone, Untangling the web: mechanisms underlying ER network formation. Biochim Biophys Acta, 2013. 1833(11): p. 2492–8.

5. Walter, P. and D. Ron, The unfolded protein response: from stress pathway to homeostatic regulation. Science, 2011. 334: p. 1081–1086.

6. Ellgaard, L. and A. Helenius, Quality control in the endoplasmic reticulum. Nature Reviews in Molecular Cell Biology, 2003. 4: p. 181–191.

7. Shao, S. and R.S. Hegde, Membrane protein insertion at the endoplasmic reticulum. Annu Rev Cell Dev Biol, 2011. 27: p. 25–56.

8. Obacz, J., et al., Regulation of tumor-stroma interactions by the unfolded protein response. FEBS J, 2019. 286(2): p. 279–296.

9. Matlack, K.E., et al., BiP acts as a molecular ratchet during posttranslational transport of prepro-alpha factor across the ER membrane. Cell, 1999. 97(5): p. 553–64.

10. Sicari, D., A. Igbaria, and E. Chevet, Control of Protein Homeostasis in the Early Secretory Pathway: Current Status and Challenges. Cells, 2019. 8(11).

11. Carvalho, P., V. Goder, and T.A. Rapoport, Distinct ubiquitin-ligase complexes define convergent pathways for the degradation of ER proteins. Cell, 2006. 126(2): p. 361–73.

12. Travers, K.J., et al., Functional and genomic analyses reveal an essential coordination between the unfolded protein response and ER-associated degradation. CELL, 2000. 101(3): p. 249–58.

13. Harding, H.P., et al., Perk is essential for translational regulation and cell survival during the unfolded protein response. Mol Cell, 2000. 5(5): p. 897–904.

14. Oakes, S.A., Endoplasmic Reticulum Stress Signaling in Cancer Cells. Am J Pathol, 2020. 190(5): p. 934–946.

15. Harding, H.P., Y. Zhang, and D. Ron, Protein translation and folding are coupled by an endoplasmic-reticulum-resident kinase. Nature, 1999. 397(6716): p. 271–4.

16. Wang, Y., et al., Activation of ATF6 and an ATF6 DNA binding site by the endoplasmic reticulum stress response. J Biol Chem, 2000. 275(35): p. 27013–20.

17. Yoshida, H., et al., XBP1 mRNA is induced by ATF6 and spliced by IRE1 in response to ER stress to produce a highly active transcription factor. Cell, 2001. 107(7): p. 881–91.

18. Calfon, M., et al., IRE1 couples endoplasmic reticulum load to secretory capacity by processing the XBP-1 mRNA. Nature, 2002. 415(6867): p. 92–6.

19. Lee, A.H., N.N. Iwakoshi, and L.H. Glimcher, XBP-1 regulates a subset of endoplasmic reticulum resident chaperone genes in the unfolded protein response. Mol Cell Biol, 2003. 23(21): p. 7448–59.

20. Hollien, J. and J.S. Weissman, Decay of endoplasmic reticulum-localized mRNAs during the unfolded protein response. Science, 2006. 313(5783): p. 104–7.

21. Kumar, R., et al., Brain ischemia and reperfusion activates the eukaryotic initiation factor 2alpha kinase, PERK. J Neurochem, 2001. 77(5): p. 1418–21.

22. Igbaria, A., et al., Chaperone-mediated reflux of secretory proteins to the cytosol during endoplasmic reticulum stress. Proc Natl Acad Sci U S A, 2019. 116(23): p. 11291–11298.

23. Sicari, D., et al., Reflux of Endoplasmic Reticulum proteins to the cytosol inactivates tumor suppressors. EMBO Rep, 2021: p. e51412.

24. Lajoie, P. and E.L. Snapp, Size-dependent secretory protein reflux into the cytosol in association with acute endoplasmic reticulum stress. Traffic, 2020.

25. Piette, B.L., et al., Comprehensive interactome profiling of the human Hsp70 network highlights functional differentiation of J domains. Mol Cell, 2021. 81(12): p. 2549–2565 e8.

26. Malinverni, D., et al., Data-driven large-scale genomic analysis reveals an intricate phylogenetic and functional landscape in J-domain proteins. Proc Natl Acad Sci U S A, 2023. 120(32): p. e2218217120.

27. Gilbert, L.A., et al., Genome-Scale CRISPR-Mediated Control of Gene Repression and Activation. Cell, 2014. 159(3): p. 647–61.

28. Fessart, D., et al., The Anterior GRadient (AGR) family proteins in epithelial ovarian cancer. J Exp Clin Cancer Res, 2021. 40(1): p. 271.

29. Fessart, D., et al., Integrative analysis of genomic and transcriptomic alterations of AGR2 and AGR3 in cancer. Open Biol, 2022. 12(7): p. 220068.

30. Goodwin, E.C., et al., BiP and multiple DNAJ molecular chaperones in the endoplasmic reticulum are required for efficient simian virus 40 infection. MBio, 2011. 2(3): p. e00101–11.

31. Holden, P. and W.A. Horton, Crude subcellular fractionation of cultured mammalian cell lines. BMC Res Notes, 2009. 2: p. 243.

32. Sicari, D., et al., A guide to assessing endoplasmic reticulum homeostasis and stress in mammalian systems. FEBS J, 2020. 287(1): p. 27–42.

