## Supplementary material for "Chaperone Complexes From The Endoplasmic Reticulum (ER) And The Cytosol Inhibit wt-p53 By Activation The ER To Cytosol Signaling"

### Supplementary Figure Legends

**Figure S1:** HLJ-1 putative orthologues share the same topology as HLJ-1. **(A)** Schematic showing the comparison between HLJ-1 topology and the five putative orthologs as described [25] with a J-domain facing the cytosol and small ER domain.

**Figure S2:** **(A-B)** Immunoblot showing the expression of DNAJB12 and DNAJB14 in A549 cells transfected with siRNA-targeted DNAJB12, DNAJB14, or both. **(C)** Subcellular protein fractionation (Digitonin fraction) of AGR2, DNAJB11, and HYOU1 in DNAJB12, DNAJB14, or DNAJB12/14 -silenced A549 cells during ER stress with tunicamycin (Tm) and quantifies in Figure 2A-C.

**Figure S3:** **(A-B)** Immunoblot showing the expression of DNAJB12 and DNAJB14 in A549 cells overexpressing either WT and QPD mutant of DNAJB12 or DNAJB14 respectively for 12hrs (left) or 24hrs (right). **(C)** cell viability of A549 cells overexpressing either WT and QPD mutant of DNAJB12 or DNAJB14 for the indicated time points. **(D)** Overexpression of either WT and QPD mutant of DNAJB12 or DNAJB14 does not induce ER stress and unfolded protein response activation, as shown by the relative mRNA levels of spliced Xbp1 and Bip.

**Supplementary Table**

**Table S1:** Alignment of the yeast HLJ-1 protein sequence against many different databases, including human, mouse, rat, zebrafish, fly, mosquito, worm, and fission yeast using the DRSC/TRiP Functional Genomics Resources & DRSC-BTRR, DIOPT version 9, Harvard medical school[Ψ].

[Ψ] Yanhui Hu, Ian Flockhart, Arunachalam Vinayagam, Clemens Bergwitz, Bonnie Berger, Norbert Perrimon, and Stephanie E Mohr. 2011. "An integrative approach to ortholog prediction for disease-focused and other functional studies." BMC Bioinformatics, 12, Pp. 357

Figure-S1

**A**

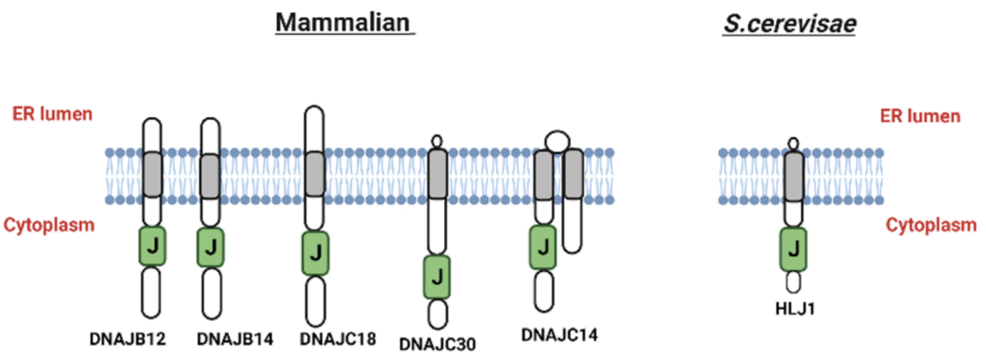


Dabsan et al, Figure S1

Figure-S2

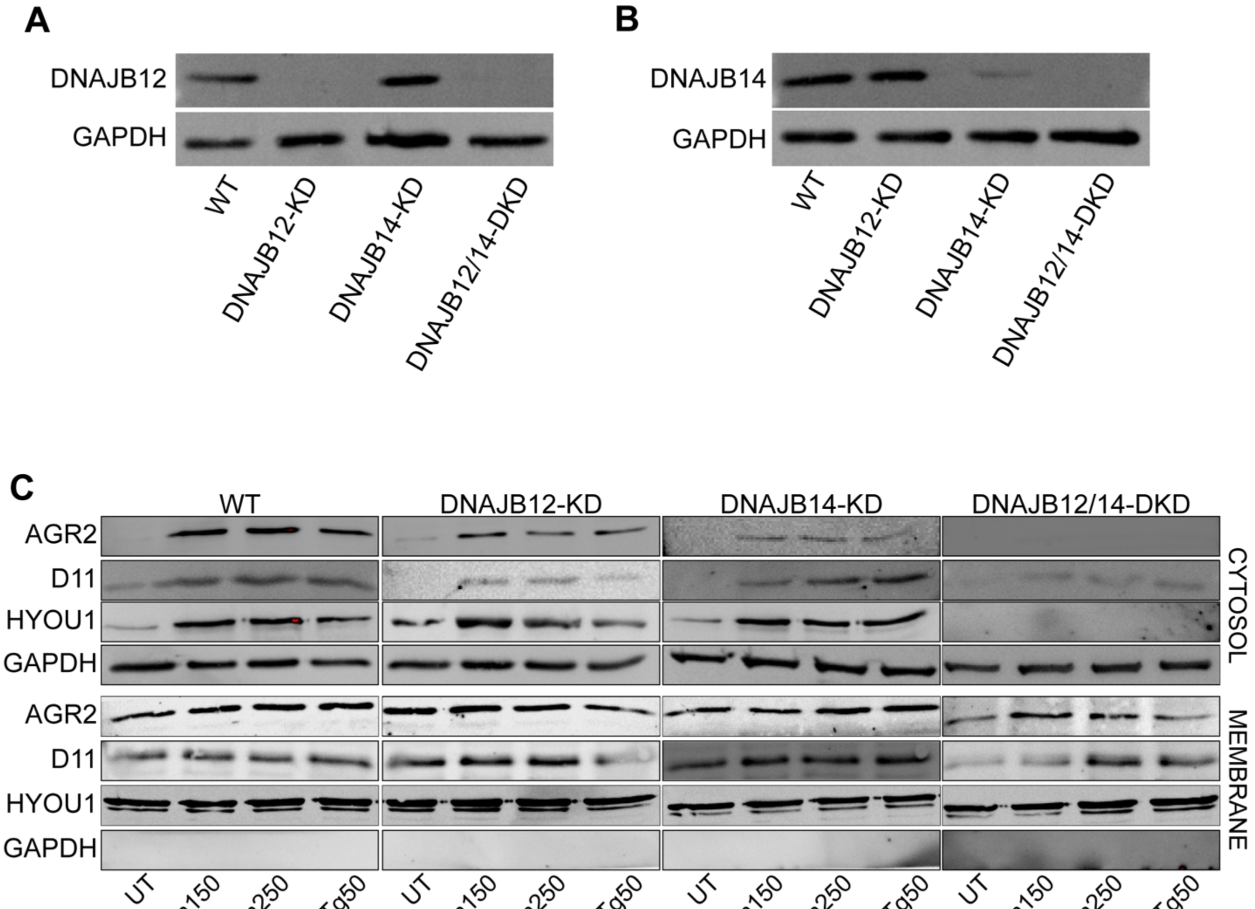


Dabsan et al, Figure S2

Figure-S3

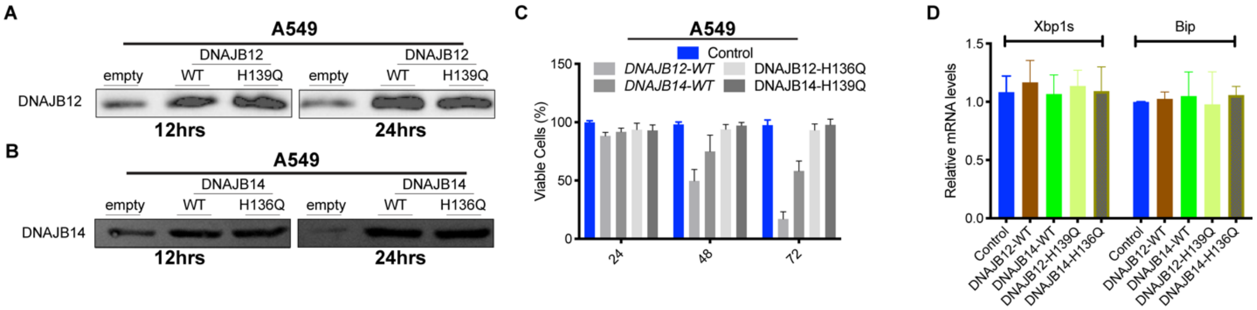


Dabsan et al, Figure S3

Table-S1

| Input Order | Search Term | Yeast GeneID | SGDID | Yeast Symbol | Species 2 | Human GeneID | Human Species Gene ID | Human Symbol | Ens |
|---|---|---|---|---|---|---|---|---|---|
| 1 | HLJ1 | 855196 | S000004771 | HLJ1 | **Human** | 54788 | 14891 | DNAJB12 | ENS |
| 1 | HLJ1 | 855196 | S000004771 | HLJ1 | **Human** | 79982 | 25881 | DNAJB14 | ENS |
| 1 | HLJ1 | 855196 | S000004771 | HLJ1 | **Human** | 202052 | 28429 | DNAJC18 | ENS |
| 1 | HLJ1 | 855196 | S000004771 | HLJ1 | **Human** | 3300 | 5228 | DNAJB2 | ENS |
| 1 | HLJ1 | 855196 | S000004771 | HLJ1 | **Human** | 10049 | 14888 | DNAJB6 | ENS |
| 1 | HLJ1 | 855196 | S000004771 | HLJ1 | **Human** | 150353 | 24986 | DNAJB7 | ENS |
| 1 | HLJ1 | 855196 | S000004771 | HLJ1 | **Human** | 165721 | 23699 | DNAJB8 | ENS |
| 1 | HLJ1 | 855196 | S000004771 | HLJ1 | **Human** | 3337 | 5270 | DNAJB1 | ENS |
| 1 | HLJ1 | 855196 | S000004771 | HLJ1 | **Human** | 4189 | 6968 | DNAJB9 | ENS |
| 1 | HLJ1 | 855196 | S000004771 | HLJ1 | **Human** | 414061 | 32397 | DNAJB3 | ENS |
| 1 | HLJ1 | 855196 | S000004771 | HLJ1 | **Human** | 85406 | 24581 | DNAJC14 | ENS |
| 1 | HLJ1 | 855196 | S000004771 | HLJ1 | **Human** | 11080 | 14886 | DNAJB4 | ENS |
| 1 | HLJ1 | 855196 | S000004771 | HLJ1 | **Human** | 25822 | 14887 | DNAJB5 | ENS |
| 1 | HLJ1 | 855196 | S000004771 | HLJ1 | **Human** | 51726 | 14889 | DNAJB11 | ENS |
| 1 | HLJ1 | 855196 | S000004771 | HLJ1 | **Human** | 10294 | 14884 | DNAJA2 | ENS |
| 1 | HLJ1 | 855196 | S000004771 | HLJ1 | **Human** | 374407 | 30718 | DNAJB13 | ENS |
| 1 | HLJ1 | 855196 | S000004771 | HLJ1 | **Human** | 134218 | 27030 | DNAJC21 | ENS |

| Input Order | Search Term | Yeast GeneID | SGDID | Yeast Symbol | Species 2 | Mouse GeneID | Mouse Species Gene ID | Mouse Symbol | Ens |
|---|---|---|---|---|---|---|---|---|---|
| 1 | HLJ1 | 855196 | S000004771 | HLJ1 | **Mouse** | 70604 | 1917854 | Dnajb14 | |
| 1 | HLJ1 | 855196 | S000004771 | HLJ1 | **Mouse** | 56709 | 1931881 | Dnajb12 | |
| 1 | HLJ1 | 855196 | S000004771 | HLJ1 | **Mouse** | 76594 | 1923844 | Dnajc18 | |
| 1 | HLJ1 | 855196 | S000004771 | HLJ1 | **Mouse** | 23950 | 1344381 | Dnajb6 | |
| 1 | HLJ1 | 855196 | S000004771 | HLJ1 | **Mouse** | 57755 | 1914012 | Dnajb7 | |
| 1 | HLJ1 | 855196 | S000004771 | HLJ1 | **Mouse** | 56691 | 1922801 | Dnajb8 | |
| 1 | HLJ1 | 855196 | S000004771 | HLJ1 | **Mouse** | 74330 | 1921580 | Dnajc14 | |
| 1 | HLJ1 | 855196 | S000004771 | HLJ1 | **Mouse** | 27362 | 1351618 | Dnajb9 | |
| 1 | HLJ1 | 855196 | S000004771 | HLJ1 | **Mouse** | 15504 | 1306822 | Dnajb3 | |
| 1 | HLJ1 | 855196 | S000004771 | HLJ1 | **Mouse** | 56812 | 1928739 | Dnajb2 | |
| 1 | HLJ1 | 855196 | S000004771 | HLJ1 | **Mouse** | 75015 | 2686498 | Samd13 | |
| 1 | HLJ1 | 855196 | S000004771 | HLJ1 | **Mouse** | 230935 | 2443386 | Dnajc11 | |
| 1 | HLJ1 | 855196 | S000004771 | HLJ1 | **Mouse** | 81489 | 1931874 | Dnajb1 | |
| 1 | HLJ1 | 855196 | S000004771 | HLJ1 | **Mouse** | 56323 | 1930018 | Dnajb5 | |
| 1 | HLJ1 | 855196 | S000004771 | HLJ1 | **Mouse** | 66861 | 1914111 | Dnajc10 | |
| 1 | HLJ1 | 855196 | S000004771 | HLJ1 | **Mouse** | 56445 | 1931882 | Dnaja2 | |
| 1 | HLJ1 | 855196 | S000004771 | HLJ1 | **Mouse** | 67035 | 1914285 | Dnajb4 | |

| Input Order | Search Term | Yeast GeneID | SGDID | Yeast Symbol | Species 2 | Rat GeneID | Rat Species Gene ID | Rat Symbol | Ens |
|---|---|---|---|---|---|---|---|---|---|
| 1 | HLJ1 | 855196 | S000004771 | HLJ1 | **Rat** | 499716 | 1591949 | Dnajb14 | |
| 1 | HLJ1 | 855196 | S000004771 | HLJ1 | **Rat** | 294513 | 1359677 | Dnajb12 | |
| 1 | HLJ1 | 855196 | S000004771 | HLJ1 | **Rat** | 291677 | 1310237 | Dnajc18 | |
| 1 | HLJ1 | 855196 | S000004771 | HLJ1 | **Rat** | 56764 | 708544 | Samd13 | |
| 1 | HLJ1 | 855196 | S000004771 | HLJ1 | **Rat** | 689593 | 1591035 | Dnajb2 | |
| 1 | HLJ1 | 855196 | S000004771 | HLJ1 | **Rat** | 680216 | 1594215 | Dnajb3 | |
| 1 | HLJ1 | 855196 | S000004771 | HLJ1 | **Rat** | 685839 | 1589047 | Dnajb7 | |
| 1 | HLJ1 | 855196 | S000004771 | HLJ1 | **Rat** | 362293 | 1308207 | Dnajb6 | |
| 1 | HLJ1 | 855196 | S000004771 | HLJ1 | **Rat** | 24908 | 3070 | Dnajb9 | |
| 1 | HLJ1 | 855196 | S000004771 | HLJ1 | **Rat** | 313811 | 1307453 | Dnajb5 | |
| 1 | HLJ1 | 855196 | S000004771 | HLJ1 | **Rat** | 114481 | 620489 | Dnajc14 | |
| 1 | HLJ1 | 855196 | S000004771 | HLJ1 | **Rat** | 500253 | 1561981 | Dnajb8 | |
| 1 | HLJ1 | 855196 | S000004771 | HLJ1 | **Rat** | 362666 | 1307731 | Dnajc11 | |
| 1 | HLJ1 | 855196 | S000004771 | HLJ1 | **Rat** | 303536 | 1303226 | Dnajc7 | |
| 1 | HLJ1 | 855196 | S000004771 | HLJ1 | **Rat** | 361384 | 1304725 | Dnajb1 | |
| 1 | HLJ1 | 855196 | S000004771 | HLJ1 | **Rat** | 295549 | 1305826 | Dnajb4 | |
| 1 | HLJ1 | 855196 | S000004771 | HLJ1 | **Rat** | 308857 | 1359131 | Dnajb13 | |
| 1 | HLJ1 | 855196 | S000004771 | HLJ1 | **Rat** | 362652 | 1359395 | Dnajc16 | |
| 1 | HLJ1 | 855196 | S000004771 | HLJ1 | **Rat** | 366567 | 1307426 | Dnajc5g | |
| 1 | HLJ1 | 855196 | S000004771 | HLJ1 | **Rat** | 311329 | 1308015 | Dnajc17 | |
| 1 | HLJ1 | 855196 | S000004771 | HLJ1 | **Rat** | 295690 | 1307813 | Dnajc10 | |
| 1 | HLJ1 | 855196 | S000004771 | HLJ1 | **Rat** | 300721 | 1310035 | Dnaja4 | |
| 1 | HLJ1 | 855196 | S000004771 | HLJ1 | **Rat** | 65028 | 620942 | Dnaja1 | |
| 1 | HLJ1 | 855196 | S000004771 | HLJ1 | **Rat** | 84026 | 71001 | Dnaja2 | |

| Input Order | Search Term | Yeast GeneID | SGDID | Yeast Symbol | Species 2 | Western clawed frog | Western clawed frog Species Gene ID | Western clawed frog | Ens |
|---|---|---|---|---|---|---|---|---|---|
| 1 | HLJ1 | 855196 | S000004771 | HLJ1 | **Western clawe** | 549342 | XB-GENE-952313 | dnajb14 | |
| 1 | HLJ1 | 855196 | S000004771 | HLJ1 | **Western clawe** | 548745 | XB-GENE-944288 | dnajb12 | |
| 1 | HLJ1 | 855196 | S000004771 | HLJ1 | **Western clawe** | 100124938 | XB-GENE-5795683 | dnajc18 | |
| 1 | HLJ1 | 855196 | S000004771 | HLJ1 | **Western clawe** | 548454 | XB-GENE-951530 | dnajb2 | |
| 1 | HLJ1 | 855196 | S000004771 | HLJ1 | **Western clawe** | 394712 | XB-GENE-972413 | dnajb6 | |
| 1 | HLJ1 | 855196 | S000004771 | HLJ1 | **Western clawe** | 780038 | XB-GENE-942676 | dnajc11 | |
| 1 | HLJ1 | 855196 | S000004771 | HLJ1 | **Western clawe** | 496977 | XB-GENE-997043 | dnajc3 | |
| 1 | HLJ1 | 855196 | S000004771 | HLJ1 | **Western clawe** | 100145451 | XB-GENE-5718163 | dnaja3 | |

| Input Order | Search Term | Yeast GeneID | SGDID | Yeast Symbol | Species 2 | Zebrafish GeneID | Zebrafish Species Gene ID | Zebrafish Symbol | Ens |
|---|---|---|---|---|---|---|---|---|---|
| 1 | HLJ1 | 855196 | S000004771 | HLJ1 | **Zebrafish** | 324005 | ZDB-GENE-030131-2725 | dnajb12a | |
| 1 | HLJ1 | 855196 | S000004771 | HLJ1 | **Zebrafish** | 792752 | ZDB-GENE-061110-138 | dnajb14 | |
| 1 | HLJ1 | 855196 | S000004771 | HLJ1 | **Zebrafish** | 100037354 | ZDB-GENE-070410-128 | dnajb12b | |
| 1 | HLJ1 | 855196 | S000004771 | HLJ1 | **Zebrafish** | 559928 | ZDB-GENE-030131-8019 | dnajc18 | |
| 1 | HLJ1 | 855196 | S000004771 | HLJ1 | **Zebrafish** | 561686 | ZDB-GENE-061013-537 | dnajb2 | |
| 1 | HLJ1 | 855196 | S000004771 | HLJ1 | **Zebrafish** | 641576 | ZDB-GENE-051127-45 | zgc:122979 | |
| 1 | HLJ1 | 855196 | S000004771 | HLJ1 | **Zebrafish** | 436626 | ZDB-GENE-040718-45 | dnajb6a | |
| 1 | HLJ1 | 855196 | S000004771 | HLJ1 | **Zebrafish** | 393275 | ZDB-GENE-040426-1122 | dnajb6b | |
| 1 | HLJ1 | 855196 | S000004771 | HLJ1 | **Zebrafish** | 101884054 | ZDB-GENE-061013-762 | zgc:152986 | |
| 1 | HLJ1 | 855196 | S000004771 | HLJ1 | **Zebrafish** | 436706 | ZDB-GENE-040718-130 | dnajc9 | |
| 1 | HLJ1 | 855196 | S000004771 | HLJ1 | **Zebrafish** | 393216 | ZDB-GENE-040426-894 | dnajc17 | |
| 1 | HLJ1 | 855196 | S000004771 | HLJ1 | **Zebrafish** | 100535820 | ZDB-GENE-121105-7 | dnajc14 | |
| 1 | HLJ1 | 855196 | S000004771 | HLJ1 | **Zebrafish** | 565734 | ZDB-GENE-131121-532 | dnajc30b | |
| 1 | HLJ1 | 855196 | S000004771 | HLJ1 | **Zebrafish** | 445061 | ZDB-GENE-040801-192 | dnajb4 | |
| 1 | HLJ1 | 855196 | S000004771 | HLJ1 | **Zebrafish** | 327244 | ZDB-GENE-030131-5455 | dnajb1b | |
| 1 | HLJ1 | 855196 | S000004771 | HLJ1 | **Zebrafish** | [illegible] | [illegible] | [illegible] | |
